# Chimeric PRMT6 protein produced by an endogenous retrovirus promoter regulates cell fate decision in mouse preimplantation embryos

**DOI:** 10.1101/2022.08.23.505038

**Authors:** Shinnosuke Honda, Maho Hatamura, Yuri Kunimoto, Shuntaro Ikeda, Naojiro Minami

## Abstract

Murine endogenous retrovirus with leucine tRNA primer, also known as MERVL, is expressed during zygotic genome activation in mammalian embryos. Here we show that protein arginine N-methyltransferase 6 (*Prmt6*) forms a chimeric transcript with MT2B2, one of the long terminal repeat sequences of MERVL, and is translated into an elongated chimeric protein (PRMT6^MT2B2^) whose function differs from that of the canonical PRMT6 protein (PRMT6^CAN^) in mouse preimplantation embryos. Overexpression of PRMT6^CAN^ in fibroblast cells increased asymmetric dimethylation of the third arginine residue of both histone H2A (H2AR3me2a) and histone H4 (H4R3me2a), while overexpression of PRMT6^MT2B2^ increased only H2AR3me2a. In addition, overexpression of PRMT6^MT2B2^ in one blastomere of mouse two-cell embryos promoted cell proliferation and differentiation of the blastomere into epiblast cells at the blastocyst stage, while overexpression of PRMT6^CAN^ repressed cell proliferation. This is the first report of the translation of a chimeric protein (PRMT6^MT2B2^) in mouse preimplantation embryos. Our results suggest that analyzing chimeric transcripts with MERVL will provide insight into the relationship between zygotic genome activation and subsequent intra- and extra-cellular lineage determination.

## Introduction

Immediately after fertilization, mammalian embryos have low transcriptional activity and translate proteins from mRNA stored in oocytes. In mouse embryos, transcription subsequently begins with two waves of zygotic genome activation (ZGA), namely minor and major ZGA, the former during the S phase of the one-cell stage and the latter during the late two-cell stage (Minami et al., 2007; Schulz and Harrison, 2019). Before and after major ZGA, the epigenome of the fertilized egg changes dynamically (Aoki, 2022; Burton and Torres-Padilla, 2010; Xu et al., 2021). Epigenetic modifications, including DNA methylation and histone modifications, are known to be important for cell differentiation during development (Wu and Sun, 2006). For example, *Carm1*, encoding a protein arginine methyltransferase, has been reported to be involved in the differentiation of blastomeres into epiblast cells through the dimethylation of histone arginine residues in mouse preimplantation embryos (Torres-Padilla et al., 2007; White et al., 2016). Endogenous retroviruses (ERVs) are one of the components of transposable elements and occupy about 10% of human and mouse genomes (Ito et al., 2020; Mouse Genome Sequencing Consortium, 2002). Although ERVs are among the earliest transcribed genetic material in mouse preimplantation embryos (Kigami et al., 2003) and various studies have shown that ERVs are expressed in mammalian preimplantation embryos (Evsikov et al., 2004; Poznanski and Calarco; Wang et al., 2001), there is little direct evidence that ERVs regulate cell differentiation. The high abundance of ERVs in the mammalian genome (Ito et al., 2020) and the fact that several ERV-derived genes, such as *Syncytin* and *Peg10*, are essential for placentation (Dupressoir et al., 2009; Dupressoir et al., 2012; Mi et al., 2000; Suzuki et al., 2007) suggest that ERVs were involved in the evolution of the placenta in mammals. Murine or human endogenous retrovirus with leucine tRNA primer (MERVL or HERVL, respectively) is a transposable element expressed during major ZGA in mice and humans (Franke et al., 2017; Kigami et al., 2003; Macfarlan et al., 2012; Wang et al., 2001). It has been reported that MERVL-activated mouse embryonic stem cells injected into eight-cell embryos can also contribute to extra-embryonic tissues (Yang et al., 2020), suggesting that some MERVL/HERVL elements may regulate the totipotency of mammalian embryos. Recently, a MERVL-derived non-coding RNA called LincGET was shown to control the differentiation of embryonic and extra-embryonic tissues in mouse preimplantation embryos (Wang et al., 2018), but the function of the vast majority of MERVL elements remains to be elucidated. Several studies have shown that the long terminal repeat of MERVL generates chimeric transcripts with host genes during early embryonic development (Macfarlan et al., 2012; Peaston et al., 2004). Because MERVL has been reported to be one of the earliest transcribed genomic sequences in preimplantation embryos (Kigami et al., 2003; Wang et al., 2001), we hypothesized that MERVL chimeric transcripts contribute to the cell lineage differentiation that occurs after major ZGA (Chazaud and Yamanaka, 2016).

Reanalysis of previously reported RNA-sequencing (RNA-seq) data revealed that protein arginine N-methyltransferase 6 (*Prmt6*) expression is activated by the MERVL promoter sequence in preimplantation embryos. Like *Carm1*, *Prmt6* encodes a Type-I protein arginine methyltransferase (Di Lorenzo and Bedford, 2011) and has been shown to regulate the epigenome by the asymmetric di-methylation of arginine residues in histone proteins (Guccione et al., 2007; Hyllus et al., 2007; Kirmizis et al., 2007). Furthermore, overexpression or suppression of *Prmt6* in embryonic stem cells results in the loss of pluripotency and in the increased expression of differentiation marker genes (Lee et al., 2012). In the course of analyzing transcripts involved in MERVL in mouse preimplantation embryos, it was discovered that *Prmt6* has two transcripts of different lengths. One is the canonical *Prmt6* transcript (*Prmt6^CAN^*), and the other is *Prmt6^MT2B2^*, defined as the chimeric transcript of *Prmt6^CAN^*with MT2B2, the MERVL long terminal repeat sequence. It was also found that the chimeric transcript produced an elongated PRMT6 chimeric protein (PRMT6^MT2B2^). In addition, the chimeric protein has different histone modification specificity from that of the canonical PRMT6 protein (PRMT6^CAN^), and overexpression of the chimeric protein in one blastomere of two-cell embryos promotes its contribution to the embryonic cell lineage. These findings shed light on a new aspect of the differentiation mechanism during embryogenesis.

## Results

### Screening of candidate MERVL chimeric transcripts associated with cell lineage differentiation

Macfarlan et al. published a list of MERVL chimeric transcripts in mouse two-cell stage embryos (Macfarlan et al., 2012), and we compared these transcripts with histone modification–related genes (Fig. 1A). Although there was no statistically significant bias, eight genes (*Atxn7*, *Brms1l*, *Ctcf*, *Kdm4c*, *Pcgf5*, *Prmt6*, *Rnf20*, and *Smyd3*) overlapped, and half of them (*Ctcf*, *Kdm4c*, *Rnf20*, and *Smyd3*) had already been reported to be important in early embryonic development (Ooga et al., 2015; Suzuki et al., 2015; Wan et al., 2008; Wang et al., 2010). Analysis of a published RNA-seq dataset (Zhang et al., 2016) revealed that the expression levels of four of the eight genes (*Ctcf*, *Kdm4c*, *Pcgf5*, and *Prmt6*) were more than twice as high in late two-cell embryos compared with early two-cell embryos (Fig. 1B)

**Fig. 1.**
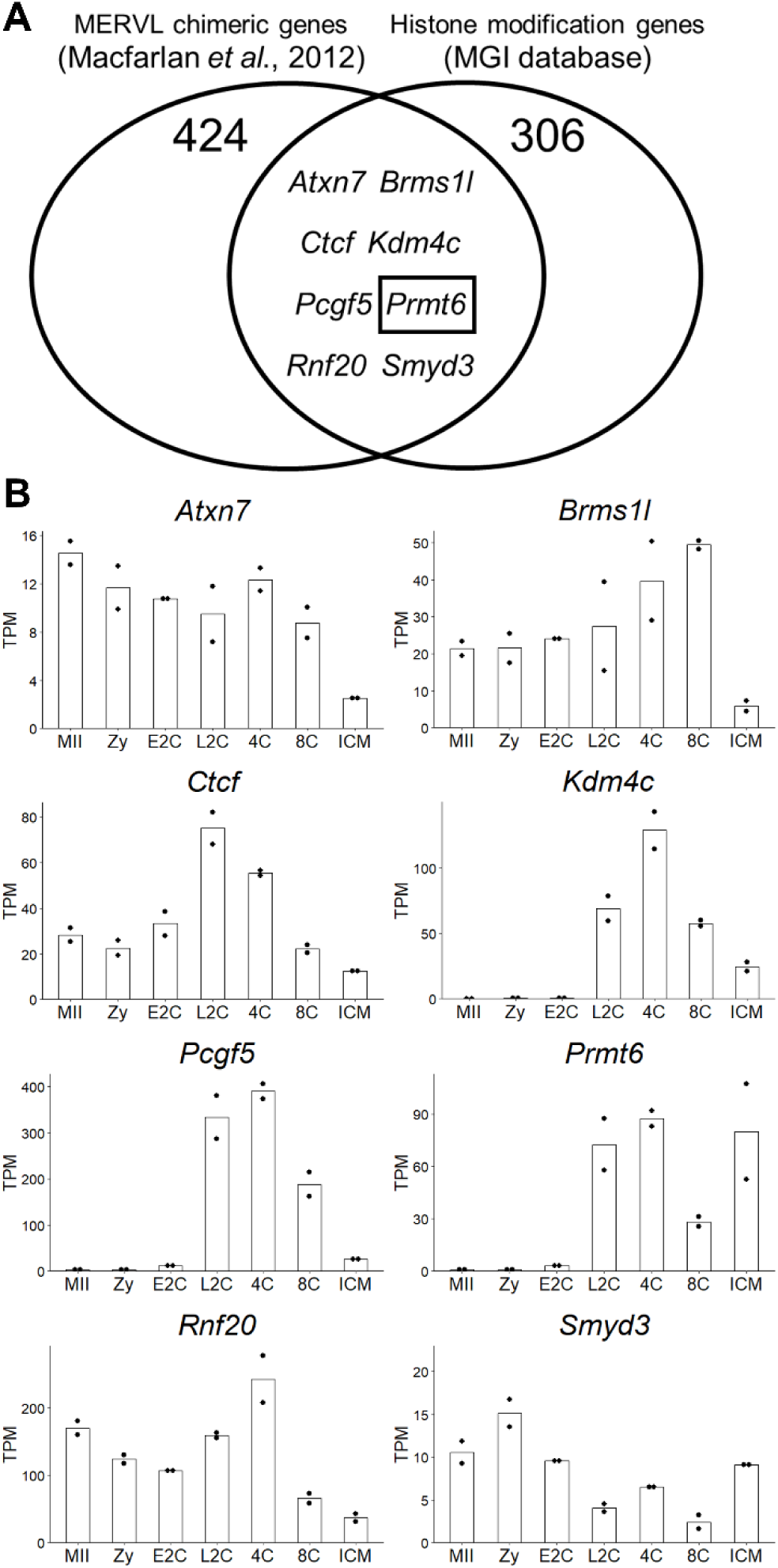
Verification of MERVL-mediated chimeric genes. (A) Venn diagram showing overlap of genes expressed as chimeric transcripts with MERVL- and histone modification–related genes. The list of MERVL chimeric genes were obtained from published data (Macfarlan et al., 2012), and the list of histone modification-related genes were obtained from MGI database (http://www.informatics.jax.org) in Jan. 2019. (B) The expression patterns of eight overlapping genes in MII oocytes and preimplantation embryos. Dots represent replicates. Data were obtained from GSE71434. MII: MII oocytes, Zy: zygotes, E2C: early two-cell embryos, 4C: four-cell embryos, 8C: eight-cell embryos, ICM: inner cell mass cells.

### Expression dynamics of chimeric and canonical Prmt6 mRNA in mouse preimplantation embryos

Genomic analysis in mice revealed that MT2B2 is located 212 bp upstream of the first exon of *Prmt6*, and that the chimeric transcript is transcribed from around 380 bp upstream of the canonical transcription start site. To determine the expression profiles of *Prmt6^MT2B2^*and *Prmt6^CAN^*, the amounts of total *Prmt6* transcripts (*Prmt6^MT2B2^* and *Prmt6^CAN^*) and *Prmt6^MT2B2^*during preimplantation development were determined by quantitative RT-PCR (Fig. 2A). As expected, the amounts of total *Prmt6* transcripts and *Prmt6^MT2B2^* transcripts in the late two-cell stage were increased at major ZGA and gradually decreased by the blastocyst stage (Fig. 2B). At the blastocyst stage, *Prmt6^MT2B2^*transcripts had almost completely disappeared, but *Prmt6^CAN^* transcripts were still present, albeit in small amounts (Fig. 2B). To compare the exact copy numbers of each transcript, absolute quantification was performed using late two-cell embryos and blastocysts. In late two-cell embryos, the amounts of total *Prmt6* transcripts and chimeric transcripts were almost the same (Fig. 2C). In blastocysts, on the other hand, the copy number of *Prmt6^MT2B2^*transcripts was significantly lower than that of total *Prmt6* transcripts (Fig. 2C). Furthermore, microinjecting zygotes with si*Prmt6^MT2B2^* (siRNA targeting the sequence between the MT2B2 and *Prmt6^CAN^* transcription start sites) or si*Prmt6^CAN^*-1 or si*Prmt6^CAN^*-2 (siRNAs targeting the *Prmt6* exon sequence, see Fig. 2A) resulted in a significant decrease in transcript levels of both total *Prmt6* transcripts and *Prmt6^MT2B2^*transcripts at the late two-cell stage (Fig. 2D). Since trimethylation of the fourth lysine residue of histone H3 (H3K4me3) is generally increased around transcription start sites, we analyzed the previously reported chromatin immunoprecipitation sequencing (ChIP-seq) data of H3K4me3 (Liu et al., 2016) in preimplantation embryos and found that the weak H3K4me3 ChIP-seq peak appeared at the MT2B2 region from the 2-cell stage and became stronger at the 4-cell stage (Fig. 2E). On the other hand, the H3K4me3 peak at just before the first exon of *Prmt6^CAN^* was observed from the four-cell stage and get higher at the eight-cell stage (Fig. 2E).

**Fig. 2.**
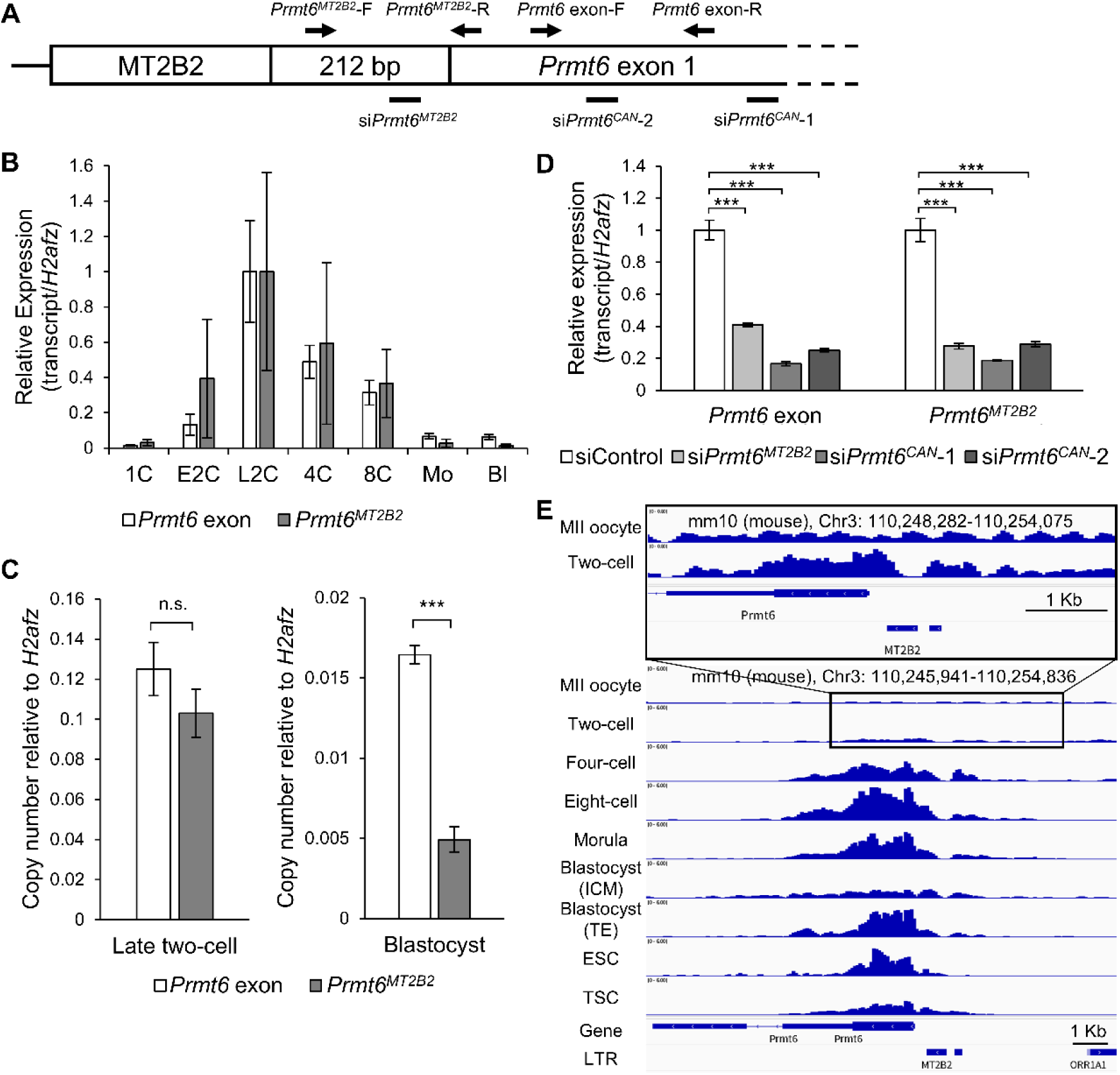
Gene structure and transcriptional profile of *Prmt6* in mice. (A) Schematic diagram showing the positions of MT2B2 and *Prmt6* in the mouse genome. MT2B2 is located 212 bp upstream from the transcription start site of *Prmt6*. The names of primers and targeting loci (arrows) are indicated above the diagram, and the names of siRNAs and their targeting sites (bars) are indicated underneath. (B) qRT-PCR analysis of the *Prmt6* exon and *Prmt6^MT2B2^* transcripts during mouse preimplantation development. The amount of mRNA in one-cell embryos was defined as 1, and the expression levels were normalized to *H2afz* as an internal control. Data are expressed as mean ± s.e.m. (n=3). (C) Comparison of expression levels of the *Prmt6* exon and *Prmt6^MT2B2^* in late two-cell embryos and blastocysts based on absolute quantification by qRT-PCR. Each graph shows the expression levels of each transcript in late two-cell embryos and blastocysts. Copy numbers were normalized to *H2afz* copy numbers. Data are expressed as mean ± s.e.m. (n=5). Student’s t-test, ***P < 0.001; n.s., not significant. (D) qRT-PCR analysis of *Prmt6* and *Prmt6^MT2B2^* mRNA abundance in embryos injected with si*Prmt6^MT2B2^*, si*Prmt6*-1, si*Prmt6*-2, or siControl in the late two-cell stage. Injection was performed 3–5 h after IVF. The amount of mRNA in siControl-injected embryos was defined as 1, and the expression levels were normalized to *H2afz* as an internal control. Data are expressed as mean ± s.e.m. (n=3). Dunnett’s test, ***P < 0.001. (E) Genome browser view of H3K4me3 ChIP-seq data from each stage of mouse preimplantation embryos. Data for MII oocytes, two-cell embryos, four-cell embryos, eight-cell embryos, morulae, inner cell mass (ICM) and trophectoderm (TE) of blastocysts, embryonic stem cells (ESC), and trophoblast stem cells (TSC) are shown. The *Prmt6^CAN^*exon and MT2B2 locations are indicated at the bottom. Data were obtained from GSE73952. A magnified view of the data for MII oocytes and two-cell embryos are shown in the upper panel. 1C: one-cell embryo, E2C: early two-cell embryo, L2C: late two-cell embryo, 4C: four-cell embryo, 8C: eight-cell embryo, Mo: morula, Bl: blastocyst

### Identification of the translation initiation codon of the PRMT6^MT2B2^ chimeric protein

Since *Prmt6^MT2B2^* has an extended sequence on the 5’ side of *Prmt6^CAN^* and two putative translation initiation codons exist in this region, we sought to determine the site where translation of the chimeric transcript began (Fig. 3A). Western blotting of the PRMT6 protein using metaphase II (MII) oocytes and fertilized eggs at each developmental stage revealed that ∼55 kDa bands were detected in oocytes and embryos during all developmental stages, and an additional ∼42 kDa band was detected only in embryos at the blastocyst stage (Fig. 3B). Since PRMT6^CAN^ has a molecular weight of ∼42 kDa, mRNAs of *Prmt6^CAN^* and *Prmt6^MT2B2^*were generated by in vitro transcription to identify the translation product of the ∼55 kDa protein detected at each developmental stage. The mRNAs transfected into NIH3T3 cells revealed that *Prmt6^MT2B2^* mRNA encodes an elongated protein of ∼55 kDa, while *Prmt6^CAN^* mRNA encodes a protein of ∼42 kDa (Fig. 3C). Consistent with the amount of transcript, PRMT6^MT2B2^ with a molecular weight of ∼55 kDa was most abundant at the late two-cell stage and gradually decreased until the blastocyst stage (Fig. 3B). There are two candidate translation initiation codons (Met1 and Met2) for *Prmt6^MT2B2^*, and Met3 is the translation initiation codon for PRMT6^CAN^ (Fig. 3A). To determine which codon *Prmt6^MT2B2^* is translated from, three types of mRNAs mutated in Met1, Met2, or Met3 (AUG to AAG) were transfected into NIH3T3 cells. Among NIH3T3 cells transfected with the three mutant mRNAs, only those with the Met1 mutant transcript failed to produce the ∼55 kDa PRMT6^MT2B2^ (Fig. 3C). The same results were obtained using mouse four-cell embryos (Fig. 3C). Bands at ∼55 kDa were detected in all four-cell embryos because these embryos express the endogenous PRMT6^MT2B2^ (Fig. 3C). Knockdown of endogenous *Prmt6* transcripts with siRNAs (si*Prmt6^CAN^*-1 or si*Prmt6^CMT2B2^*) at the one-cell stage decreased the amount of the ∼55 kDa protein at the four-cell stage (Fig. 3D).

**Fig. 3.**
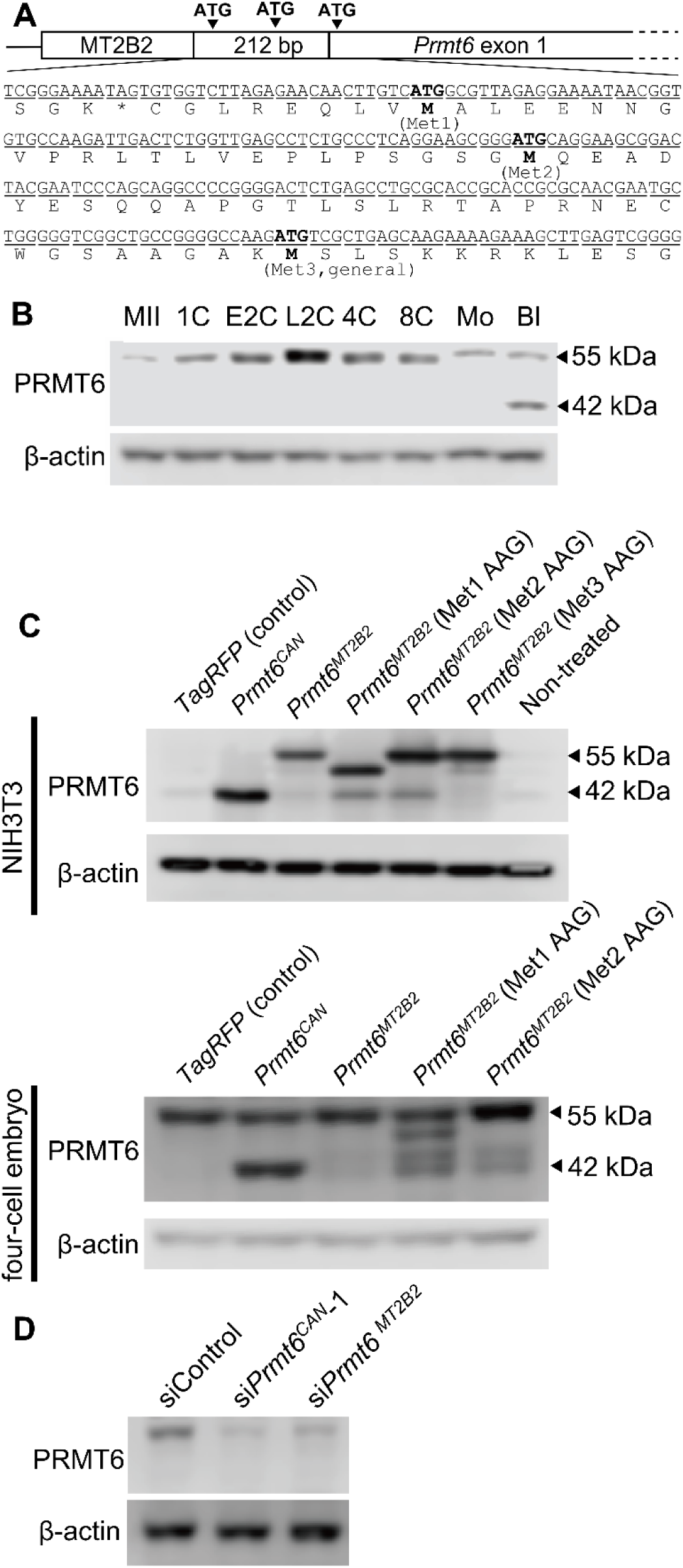
Protein expression of Prmt6. (A) Schematic diagram of the genome structure (upper) and genome and protein sequences (lower) of *Prmt6* exon 1. Candidate translation initiation codons (Met1 and Met2) and the canonical translation initiation codon (Met3) are indicated above the schematic diagram and are shown in bold in the sequences. (B) Western blotting of PRMT6 in mouse oocytes and preimplantation embryos. β-actin was used as a loading control. Fifty oocytes or embryos were used in each lane except blastocysts. Twenty-five blastocysts were used. The approximate molecular weight is on the right. The representative result is shown from three independent experiments. (C) Western blotting analysis for PRMT6 in NIH3T3 cells and four-cell embryos. NIH3T3 cells were transfected with TagRFP mRNA, *Prmt6^CAN^* mRNA, *Prmt6^MT2B2^* mRNA, *Prmt6^MT2B2^* with Met1 mutant mRNA (Met1 AAG), *Prmt6^MT2B2^* with Met2 mutant mRNA (Met2 AAG), or *Prmt6^MT2B2^* with Met3 mutant mRNA (Met3 AAG). Zygotes were injected with the same mRNAs except *Prmt6^MT2B2^* with Met3 mutant mRNA (Met3 AAG) and cultured to the four-cell stage. Fifty embryos were used in each lane. β-actin was used as a loading control. The approximate molecular weight is on the right. The representative result is shown from three independent experiments. (D) Western blotting analysis of PRMT6 protein in Prmt6-suppressed four-cell embryos. si*Prmt6^CAN^*-1, si*Prmt6^MT2B2^*, or siControl was injected into zygote cytoplasm. β-actin was used as a loading control. Fifty embryos were used for each lane. The approximate molecular weight is on the right. The representative result is shown from three independent experiments. MII: metaphase II oocyte, 1C: one-cell embryo, E2C: early two-cell embryo, L2C: late two-cell embryo, 4C: four-cell embryo, 8C: eight-cell embryo, Mo: morula, Bl: blastocyst

### Functional analysis of the PRMT6^MT2B2^ chimeric protein

Three-dimensional structure prediction using AlphaFold (Jumper et al., 2021) showed no structural change in the S-adenosylmethionine–dependent methyltransferase domain of PRMT6^MT2B2^ compared to PRMT6^CAN^ (Fig. 4A). Therefore, the ∼55 kDa PRMT6^MT2B2^ was considered to have arginine methylation activity, and the function of this protein and its involvement in cell differentiation in mammalian preimplantation embryos were examined. Since PRMT6^CAN^ is known to be a type I protein arginine methyltransferase (Di Lorenzo and Bedford, 2011), western blotting was performed using various histone arginine asymmetric dimethylation antibodies to confirm the histone arginine methylation status of NIH3T3 cells overexpressing PRMT6^CAN^ or PRMT6^MT2B2^. Overexpression of PRMT6^CAN^ significantly increased asymmetric dimethylation of the third arginine residue of both histone H2A (H2AR3me2a) and histone H4 (H4R3me2a) (Fig. 4B, 4C, 4D). By contrast, overexpression of PRMT6^MT2B2^ significantly increased only H2AR3me2a, while the amount is lower than overexpression of PRMT6^CAN^ (Fig. 4B, 4C, 4D; P<0.01).

**Fig. 4.**
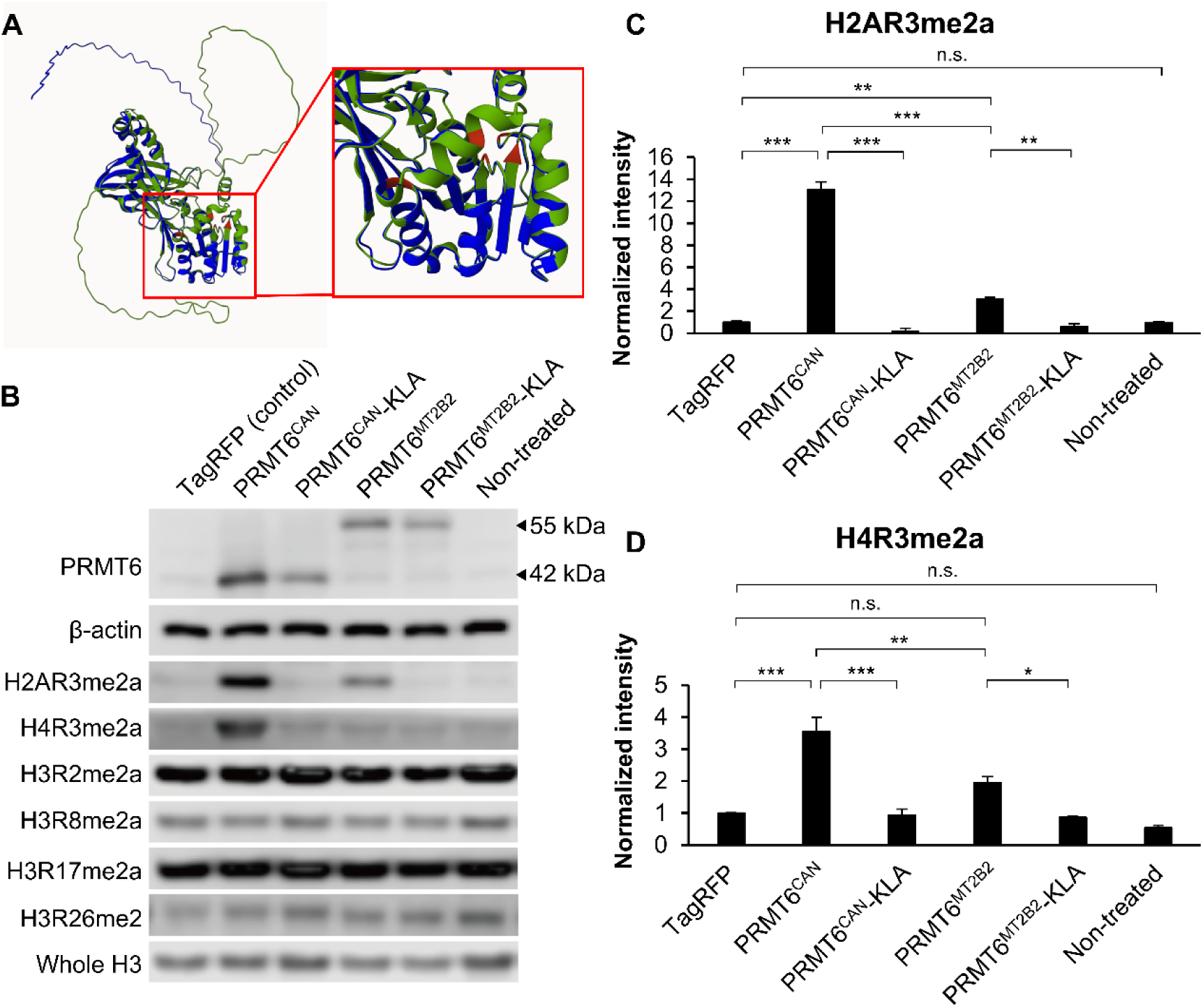
Substrate-specific histone arginine methylation activity of PRMT6^MT2B2^. (A) The 3D structures of PRMT6^CAN^ and PRMT6^MT2B2^ as predicted by AlphaFold2. The blue structure represents PRMT6^CAN^ and the green structure represents PRMT6^MT2B2^. The five S-adenosylmethionine binding sites are shown in red. The right panel shows a magnified view of the catalytic domain. (B) Western blotting analysis of various histone arginine methylation modifications in NIH3T3 cells overexpressing TagRFP, PRMT6^CAN^, PRMT6^CAN^-KLA, PRMT6^MT2B2^, or PRMT6^MT2B2^-KLA. β-actin and whole H3 were used as loading controls. KLA-containing proteins are those in which the enzyme activity is eliminated by replacing amino acids VLD with KLA. (C, D) The amounts of H2AR3me2 (C) and H4R3me2 (D) based on the intensity of (B). Overexpression of PRMT6^CAN^ increases H2AR3me2a and H4R3me2a intensity, while overexpression of PRMT6^MT2B2^ increases only H2AR3me2a intensity. Data are expressed as mean ± s.d. (n=3). Tukey-Kramer test, ***P < 0.001; **P < 0.01; *P < 0.05; n.s., not significant.

### PRMT6^MT2B2^ regulates cell differentiation in mouse preimplantation embryos

Analysis of published single cell RNA-seq data (Goolam et al., 2016) showed that *Prmt6* is asymmetrically distributed among blastomeres in four- and eight-cell embryos (Supplemental Fig. 1A). Next, we confirmed H2AR3me2a distribution among blastomeres at each developmental stage. Immunostaining of H2AR3me2a showed an asymmetrical distribution among blastomeres after the eight-cell stage and a greater abundance in the trophectoderm (TE) than in the inner cell mass (ICM) at the blastocyst stage (Supplemental Fig. 1B). *Prmt6^CAN^* or *Prmt6^MT2B2^* mRNA together with *H2B-EGFP* mRNA as a lineage tracer was microinjected into one blastomere of two-cell embryos to examine the effects on cell lineage specification in mouse preimplantation embryos (Fig. 5A). Overexpression of PRMT6^MT2B2^ significantly increased the number of H2B-EGFP– positive cells in blastocysts at 96 h post insemination (hpi) compared to overexpression of H2B-EGFP alone, while PRMT6^CAN^ overexpression reduced the number of H2B-EGFP–positive cells in blastocysts at 96 hpi (Fig. 5B, 5C). In addition, overexpression of PRMT6^MT2B2^ significantly increased the percentage of NANOG-positive epiblast cells in H2B-EGFP–positive cells in blastocysts than overexpression of H2B-EGFP alone (Fig. 5B, 5D), while this percentage was unaffected by PRMT6^CAN^ overexpression (Fig. 5D). The differentiation into trophectoderm (CDX2-positive cells) and primitive endoderm (NANOG-negative and CDX2-negative cells) were not changed by PRMT6^CAN^ or PRMT6^MT2B2^ overexpression (Supplementary Table 3). The si*Prmt6^CAN^*-1 injection into one blastomere of two-cell embryos resulted in the low number of H2B-EGFP–positive cells in blastocyst, while neither si*Prmt6^CAN^*-1 nor si*Prmt6^MT2B2^* injection influence cell differentiation (Supplementary Table 4). When *Prmt6^MT2B2^* or *Prmt6^CAN^* mRNA was injected in one-cell embryos, cell numbers were not changed comparing with EGFP mRNA injected control embryos, while the percentage of CDX2-positive cells was decreased in PRMT6^MT2B2^ overexpressed embryos (Supplementary Table 5).

**Fig. 5.**
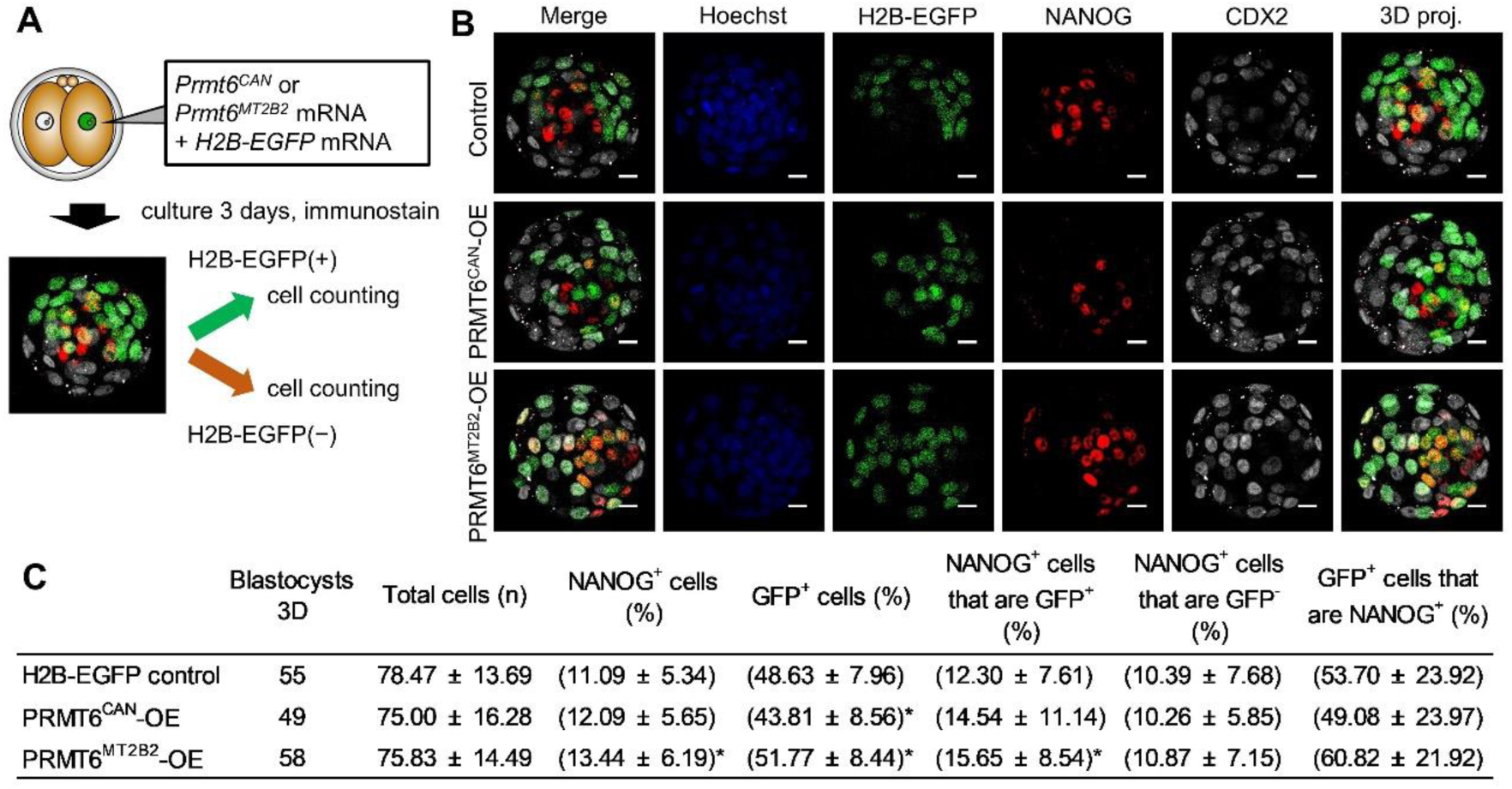
PRMT6^MT2B2^ determines the cell fate of mouse preimplantation embryos. (A) Schematic view of the cell fate contribution assay. *Prmt6^CAN^* or *Prmt6^MT2B2^* mRNA together with *H2B-EGFP* mRNA was microinjected into one blastomere of two-cell embryos. Embryos cultured until the blastocyst stage (96 hpi) were subjected to immunostaining using antibodies against NANOG (red) and CDX2 (white). Cell nuclei were stained with Hoechst (blue). The number of NANOG-positive cells was counted separately for H2B-EGFP–positive and –negative cells. (B) Immunostaining of NANOG and CDX2 in blastocysts overexpressing H2B-EGFP (control), PRMT6^CAN^ with H2B-EGFP (PRMT6^CAN^-OE), or PRMT6^MT2B2^ with H2B-EGFP (PRMT6^MT2B2^-OE). Photos are representative of immunostained blastocysts. Hoechst (blue), H2B-EGFP (marker, green), NANOG (red), CDX2 (white), and 3D maximum projections of merged images are shown. Scale bars, 20 μm. (C) Analysis of the distribution of progeny of injected blastomere at the blastocyst stage based on (B). Data were represented as mean ±s.d. Wilcoxon signed-rank test with holm’s adjustment were used for statistical analysis, *P < 0.05. Seven experimental replicates were performed. Key to table headings: Blastocysts 3D is the number of blastocysts that were all analyzed, Total cells (n) is the total number of cells in the blastocyst, NANOG^+^ cells (%) is the percentage of NANOG-positive cells out of the total number of cells in the blastocyst, GFP^+^ cells (%) is the percentage of GFP-positive cells out of the total number of cells in the blastocyst, NANOG^+^ cells that are GFP^+^ (%) is the percentage of NANOG-positive cells out of the total number of GFP-positive cells in the blastocyst, NANOG^+^ cells that are GFP^-^ (%) is the percentage of NANOG-positive cells out of the total number of GFP-negative cells in the blastocyst, and GFP^+^ cells that are NANOG^+^ (%) is the percentage of GFP-positive cells out of the total number of NANOG-positive cells in the blastocyst.

## Discussion

Among the histone modification–related genes expressed as chimeric transcripts with MERVL, the expression of four genes (*Ctcf*, *Kdm4c*, *Pcgf5*, and *Prmt6*) dramatically increased during major ZGA, suggesting that many genes expressed during this period may be mainly under control of the MERVL promoter. Indeed, the results of absolute quantitative RT-PCR (Fig. 2C) and siRNA knockdown experiments targeting the intermediate sequence between MT2B2 and *Prmt6* exon (Fig. 2D) showed that almost all *Prmt6* transcripts were *Prmt6^MT2B2^*at the late two-cell stage. In contrast, most *Prmt6^CAN^* transcripts were expressed at the blastocyst stage, indicating that the transcriptional machinery of *Prmt6* is dramatically altered during early embryogenesis. Analysis of H3K4me3 ChIP-seq data supported the dynamic change of promoter activity, but the H3K4me3 peak at the MT2B2 region at the two-cell stage is weaker than the four- and eight-cell stage. The temporal mismatch between H3K4me3 ChIP-seq data and our quantitative RT-PCR data could be due to the difference of mice strain (Bertozzi et al., 2020) or collection time of 2-cell embryos.

Overexpression experiments of *Prmt6^MT2B2^* mRNA with a mutation in the translation initiation codon revealed that the *Prmt6^MT2B2^*chimeric transcript is translated into the PRMT6^MT2B2^ chimeric protein from the 5’-extended uppermost ATG codon (Fig. 3C, 3D). PRMT6^MT2B2^ is the first MERVL chimeric protein translated from a chimeric transcript during mammalian development. Outside of early embryogenesis, one case of a MERVL chimeric transcript translated into a longer protein was reported in cancer cells (Lock et al., 2014). Since the extended 56-amino-acid sequence of PRMT6^MT2B2^ showed no homology to retrovirus-derived proteins such as the GAG protein (data not shown), further studies on the origin of the elongated sequence are needed to understand the biological functions of PRMT6^MT2B2^ chimeric proteins. Although many chimeric transcripts are expressed in cancer cells and embryonic stem cells (Babaian and Mager, 2016; Macfarlan et al., 2012), their exact translation initiation codons and biological functions remain unknown, and more comprehensive proteomic analysis is needed to understand the precise function of chimeric transcripts, including those expressed in preimplantation embryos.

Our results also revealed that the PRMT6^MT2B2^ chimeric protein has more substrate-specific and/or lower histone arginine methylation activity than PRMT6^CAN^ (Fig. 4B, 4C), suggesting that changes in epigenomic modifications may regulate developmental progression and gene expression in early preimplantation embryos. H2AR3me2a is asymmetrically distributed among blastomeres after the 8-cell stage and is more abundant in trophectoderm at the blastocyst stage (Supplemental Fig. 1B). Since PRMT6^CAN^ has a stronger ability to modify H2AR3me2a than PRMT6^MT2B2^ (Fig. 4C), it is possible that the transcription shifts from *Prmt6^MT2B2^* to *Prmt6^CAN^* in some blastomeres after the 8-cell stage, resulting in increased amounts of H2AR3me2a and enhanced differentiation to trophectoderms. PRMT6 is known to be expressed in several cancer tissues, including colorectal, breast, bladder, and lung adenocarcinomas (Avasarala et al., 2020; Lim et al., 2018; Tang et al., 2020; Veland et al., 2017; Yoshimatsu et al., 2011). Investigating how PRMT6^MT2B2^ is associated with cancer progression will provide insight into the involvement of retroviruses in cancer development (Attermann et al., 2018; Babaian and Mager, 2016; Kassiotis and Stoye, 2016; María et al., 2016) and a new perspective on cancer research. The PRMT6^CAN^ protein contains a nuclear localization signal (NLS) at the N-terminus (Fig. 3A) (Frankel et al., 2002; Herrmann et al., 2005). Since it is known that the modification of surrounding amino acids and protein conformation affect NLS function (Sorokin et al., 2007), it is possible that the extended amino acid sequence of PRMT6^MT2B2^ leads to loss of NLS function and lower catalytic activity for histone proteins. Since other arginine methyltransferases are known to methylate proteins localized in the cytoplasm (Di Lorenzo and Bedford, 2011; Litt et al., 2009), analysis of the subcellular localization of PRMT6^MT2B2^ could provide new insights into its function.

In blastocysts, most *Prmt6* transcripts are presumably *Prmt6^CAN^*based on absolute quantification (Fig. 2C). Blastomeres overexpressing PRMT6^MT2B2^ were more likely to consist of epiblast cells than non-injected blastomeres, while those overexpressing PRMT6^CAN^ did not contribute to each cell lineage (Fig. 4G). In addition, *Prmt6^MT2B2^*mRNA microinjection to one-cell embryos showed lower CDX2-positive cell rate at the blastocyst stage, which is stronger phenotype than *Prmt6^CAN^*mRNA injected embryos. These indicates that PRMT6^MT2B2^ and PRMT6^CAN^ have different functions in cell differentiation during embryogenesis. It has been reported that H3R2me2a mediated by PRMT6^CAN^ recruits Aurora B to chromosome arms (Kim et al., 2020) and that overexpression of Aurora B facilitates cell contribution to the placental cell lineage by regulating the nuclear dynamics of OCT4 in mouse preimplantation embryos (Li et al., 2017; Plachta et al., 2011). Based on these results, it is conceivable that an increased ratio of PRMT6^CAN^ to PRMT6^MT2B2^ promotes cell differentiation during the blastocyst stage through localization of PRMT6^CAN^ in the nucleus. In addition, several protein arginine methyltransferases have been reported to act redundantly (Cheng et al., 2020; Wei et al., 2021). *Carm1* was found to promote differentiation of blastomeres to epiblast cells via methylation of H3R26me2 (Goolam et al., 2016; Torres-Padilla et al., 2007; White et al., 2016), and *Prmt1* enzymic inhibitor promoted the differentiation of embryonic stem cells into primitive endoderm via Klf4 methylation (Zuo et al., 2022). These results support the possibility that PRMT6^MT2B2^, but not PRMT6^CAN^, may regulate the differentiation pathways in blastocysts.

Microinjection of PRMT6^CAN^ or si*Prmt6^CAN^*-1 in a single blastomere in two-cell stage embryos resulted in delayed cell proliferation (Fig. 4F), probably since PRMT6^CAN^ controls the cell cycle, as reported in previous studies (Kim et al., 2020; Schneider et al., 2021). On the other hand, overexpression of PRMT6^MT2B2^ promoted cell proliferation (Fig. 4F), indicating that PRMT6^CAN^ and PRMT6^MT2B2^ affect cell proliferation in different ways. Modzelewski et al. reported that the MERVL-*Cdk2ap1* chimeric transcript is translated into a shorter protein (CDK2AP1^ΔN^) than the canonical protein (CDK1AP1^CAN^) in early mammalian embryos, and that CDK2AP1^ΔN^ promotes cell proliferation while CDK2AP1^CAN^ inhibits the cell cycle (Modzelewski et al., 2021). In mice, ZGA is known to occur earlier and to have a shorter implantation period than in other mammals (Modzelewski et al., 2021). Adaptive evolution for rapid development of mouse embryos may have occurred through the insertion of MT2B2 just before the Prmt6 locus, which allowed translation of the Prmt6MT2B2 chimeric protein and may have promoted cell proliferation after ZGA. These findings and our own indicate that chimeric transcripts expressed during major ZGA mediate different functions than canonical transcripts.

In conclusion, we have shown that the chimeric transcript *Prmt6^MT2B2^*is expressed in mouse preimplantation embryos and that it generates the chimeric protein PRMT6^MT2B2^ during major ZGA. Furthermore, PRMT6^MT2B2^ expressed in early preimplantation embryos is involved in determining cell differentiation during the blastocyst stage. This study reveals that MERVL expressed at major ZGA regulates cell differentiation during embryogenesis by regulating the expression of nearby genes and the translation of their corresponding mRNA.

## Materials and Methods

### In vitro fertilization and embryo culture

Eight- to twelve-week-old ICR female mice (Japan SLC, Hamamatsu, Japan) were superovulated by injection of 7.5 IU of equine chorionic gonadotropin (eCG; ASKA Pharmaceutical, Tokyo, Japan) followed by 7.5 IU of human chorionic gonadotropin (hCG; ASKA Pharmaceutical) 48 h later. Cumulus oocyte complexes were harvested 14 h after hCG injection and placed in human tubal fluid (HTF) medium supplemented with 4 mg/ml bovine serum albumin (BSA, A3311; Sigma-Aldrich, St. Louis, MO) covered with paraffin oil (Nacalai Tesque, Kyoto, Japan). Spermatozoa were collected from the cauda epididymis of 13–20-week-old ICR male mice (Japan SLC) and cultured in HTF medium for 1 h. After capacitation, sperm were introduced into oocyte-containing droplets at a final concentration of 1 × 10^6^ cells/ml. After 3-h incubation at 37°C in an atmosphere of 5% CO_2s_, fertilized oocytes were washed with K^+^-modified simplex optimized medium (KSOM) supplemented with amino acids (Ho et al., 1995) and 1 mg/ml BSA, and then used for microinjection or cultured in the same medium under paraffin oil at 37°C in an atmosphere of 5% CO_2_ to the following stages: one-cell (12 h post insemination (hpi)), early two-cell (24 hpi), late two-cell (36 hpi), four-cell (48 hpi), eight-cell (54 hpi), morula (72 hpi), and blastocyst (96 hpi). MII oocytes were collected from cumulus oocyte complexes followed by treatment with 1% hyaluronidase to remove cumulus cells.

### Plasmid construction, in vitro transcription, and microinjection

Based on the transcription start site of *Prmt6^CAN^* predicted from the NCBI database (NM_178891.5) and that of *Prmt6^MT2B2^* predicted from previously reported RNA-seq data (Liu et al., 2016), the sequences of *Prmt6^CAN^* and *Prmt6^MT2B2^* were cloned by PCR from the cDNA of late two-cell embryos. The amplified products were digested with EcoRI and XbaI, and the fragments were incorporated into a pBluescript II SK (−) vector plasmid (Agilent Technologies Japan, Hachioji, Japan). Start codon mutant constructs were generated by mutation PCR using the PrimeSTAR Mutagenesis Basal Kit (Takara Bio, Kusatsu, Japan) by replacing AUG with AAG at Met1, Met2, or Met3. The AAG codon was chosen because this mutation has been reported to cause complete loss of translational activity in eukaryotic cells compared to other single-nucleotide mutations (Wei et al., 2013). Methylase-inactive KLA mutant constructs were generated using the same method by replacing amino acids Val–Leu–Asp at positions 89–91 in PRMT6^CAN^ with Lys–Leu–Ala to produce PRMT^CAN^-KLA (Boulanger et al., 2005), or by replacing these amino acids at positions 145–147 in PRMT6^MT2B2^ with Lys–Leu–Ala to produce PRMT6^MT2B2^-KLA. Each mRNA was synthesized with the mMESSAGE mMECHINE T7 Ultra Kit (Thermo Fisher Scientific, Waltham, MA), purified with the RNeasy Mini Kit (Qiagen, Hilden, Germany), and then dissolved in nuclease-free water. For knockdown experiments, siRNAs (RNAi, Tokyo, Japan) dissolved in nuclease-free water at a concentration of 100 µM were used for microinjection. For overexpression experiments, *Prmt6^CAN^*or *Prmt6^MT2B2^* mRNA was diluted to 250 ng/µl and co-injected with *H2B-EGFP* mRNA at a concentration of 100 ng/µl. Approximately 5–10 pl of mRNA or siRNA was microinjected into the cytoplasm of one-cell embryos at 3–5 hpi or into the cytoplasm of one blastomere of two-cell embryos at 24–26 hpi in HEPES-buffered KSOM. Microinjection was performed under an inverted microscope (IX73; Olympus, Tokyo, Japan) equipped with a piezo injector (PMAS-CT150; PRIME TECH, Tokyo, Japan) and a micromanipulator (IM-11-2; Narishige, Tokyo, Japan). The primers and siRNA sequences are listed in Supplementary Tables 1 and 2.

### Cell culture and mRNA transfection

NIH/3T3 cells were cultured in Dulbecco’s modified Eagle’s medium supplemented with 10% calf serum, 75 µg/ml penicillin, and 50 µg/ml streptomycin. mRNAs transcribed in vitro were transfected into 50–60% confluent cells using Lipofectamine MessengerMAX Transfection Reagent (Thermo Fisher Scientific). Twenty-four hours after transfection, the cells were washed in phosphate-buffered saline (PBS) once and then lysed in 1X Laemmli sample buffer.

### RNA extraction and quantitative RT-PCR

RNA extraction and reverse transcription were performed using the SuperPrep II Cell Lysis & RT Kit for qPCR (Toyobo, Osaka, Japan). Briefly, five embryos or oocytes were washed three times in PBS containing 0.5% polyvinylpyrrolidone (PVP K-30; Nacalai Tesque) (PVP-PBS) and collected in PCR tubes along with 0.5 µl of PVP-PBS. After adding 3.5 µl of lysate, all lysates were subjected to reverse transcription. Transcription levels were measured as previously described (Shikata et al., 2020) using the 2^−ΔΔCt^ method normalized against the corresponding *H2afz* levels (Jeong et al., 2014; Livak and Schmittgen, 2001). Absolute expression was determined using serial dilutions of the chimeric *Prmt6* cloned plasmid for in vitro transcription or using the pTAC plasmid in which an *H2afz* PCR fragment was cloned. The primers used for quantification are listed in Supplementary Table 2.

### Western blotting

Fifty embryos or oocytes, excluding blastocysts, were washed three times in PVP-PBS and collected in 1X Laemmli sample buffer. Twenty-five blastocysts were collected. After boiling at 95°C for 5 min, the samples were kept at −80°C until use. NIH3T3 cells were directly lysed in 1X Laemmli sample buffer. To analyze histone arginine methylation modifications in NIH3T3 cells, TagRFP, PRMT6^CAN^-KLA, or PRMT6^MT2B2^-KLA were used as negative controls for overexpression. Total protein was separated by SDS–polyacrylamide gel electrophoresis (SDS-PAGE) and transferred into a 0.22-µm-pore polyvinylidene fluoride (PVDF) membrane (Cytiva, Tokyo, Japan). After one wash in PBS containing 0.1% Tween (PBST), the membrane was blocked in PBST containing 5% BSA for 1 h at room temperature, followed by incubation with a rabbit anti-PRMT6 antibody (1:1500 dilution; 15395-1-AP; Proteintech, Rosemont, IL), a rabbit anti-H2AR3me2a antibody (1:1000 dilution; ab21574; Abcam, Cambridge, UK), a rabbit anti-H4R3me2a antibody (1:500 dilution; 39706; Active Motif, Carlsbad, CA), a rabbit anti-H3R2me2a antibody (1:1000 dilution; ab175007; Abcam), a rabbit anti-H3R8me2a antibody (1:500 dilution; 39652; Active Motif), a rabbit anti-H3R17me2a antibody (1:2000 dilution; 39710; Active Motif), a rabbit anti-H3R26me2 antibody (1:8000 dilution; 07-215; Merck, Darmstadt, Germany), a mouse anti–β-actin antibody (1:5000 dilution; A5441; Merck), or a rabbit anti–whole-H3 antibody (1:1000 dilution; #9715; Cell Signaling, Danvers, MA) in PBST containing 1% BSA overnight at 4°C. After three washes in PBST, the membrane was incubated with a horseradish peroxidase–conjugated anti-mouse secondary antibody (1:10000 dilution; NA931; Cytiva) or anti-rabbit secondary antibody (1:10000 dilution; NA934; Cytiva) in PBST containing 1% BSA for 1 h at room temperature. After three washes in PBST, the membrane was developed using ImmunoStar LD (FUJIFILM Wako Pure Chemical Corporation, Osaka, Japan) and scanned with c-DiGit (LI-COR, Lincoln, NE). Densitometric quantification of the immunoblot bands was performed using Image Studio software (LI-COR).

### Immunofluorescence

Embryos were fixed in PBS containing 4% paraformaldehyde for 20 min at 28°C. After three washes in PVP-PBS, embryos were permeabilized with 0.5% Triton X-100 (Sigma-Aldrich) in PBS for 40 min at 28°C. After washing three times in PVP-PBS, embryos were blocked in PBS containing 1.5% BSA, 0.02% Tween 20, and 0.2% sodium azide (blocking buffer) for 1 h at 28°C, and then incubated overnight at 4°C with a rabbit anti-NANOG antibody (1:100 dilution; RCAB002P-F; ReproCELL, Kanagawa, Japan) and a mouse anti-CDX2 antibody (1:100 dilution; MU392A-UC; Biogenex, Fremont, CA). After three washes in blocking buffer, embryos were incubated in blocking buffer containing secondary antibodies (Alexa Fluor 555 donkey anti-rabbit IgG, 1:500 dilution, A31572, Thermo Fisher Scientific; Alexa Fluor 647 donkey anti-mouse IgG, 1:500 dilution, A31571, Thermo Fisher Scientific) for 1 h at 28°C. After washing three times in blocking buffer, nuclei were stained in blocking buffer containing 10 µg/ml Hoechst 33342 (Sigma-Aldrich) for 10 min at 28°C. Stained embryos were mounted on slide glass in blocking buffer containing 50% glycerol and signals were observed with a confocal fluorescent microscope (LSM880; Zeiss, Oberkochen, Germany). H2B-EGFP protein was detected based on its own fluorescence.

### RNA-seq and ChIP-seq data processing

The bulk RNA-seq data (GSE71434) and the single cell RNA-seq data (E-MTAB-3321) of embryos of each stage were downloaded and then mapped to the mm10 with STAR (v. 2.7.10a) (Dobin et al., 2013). The gene expression level was estimated using RSEM (v. 1.3.3) (Li and Dewey, 2011) and normalized with the transcripts per kilobase million method. The H3K4me3 ChIP-seq data of mouse preimplantation embryos were downloaded from GSE73952 and mapped to the mm10 with bowtie2 (v. 2.4.5) (Langmead and Salzberg, 2012). In all cases, Bam files of biological replicates were merged using samtools (v. 1.10) (Li et al., 2009), normalized by counts per million using deeptools (v. 3.5.1) (Ramírez et al., 2016), and visualized using Integrative Genomics Viewer software (v. 2.12.1) (Robinson et al., 2011).

### Three-dimensional Structure Prediction of the PRMT6^MT2B2^ Protein

PRMT6^CAN^ and PRMT6^MT2B2^ protein structure prediction was performed using Alphafold (Jumper et al., 2021) on the Google Colaboratory site (https://colab.research.google.com/github/sokrypton/ColabFold/blob/main/AlphaFold2.i pynb). Obtained data were visualized using the Mol* site (Sehnal et al., 2021) (https://molstar.org/).

### Statistical analyses

Quantitative RT-PCR data were analyzed by Student’s t-test for pairwise comparisons or by Dunnett’s test for multiple comparisons. Quantification of western blotting was performed using the Tukey-Kramer test. Cell contribution assays were analyzed with the Wilcoxon signed-rank test with holm’s adjustment. All analyses were performed using R (v. 4.2.1), and significance was accepted at P-values < 0.05.

### Ethical approval for the use of animals

All animal experiments were approved by the Animal Research Committee of Kyoto University (Permit Numbers: R3-17) and were performed in accordance with the committee’s guidelines.

## Acknowledgments

Confocal fluorescent microscopy was performed at the iCeMS Analysis Center, Institute for Integrated Cell-Material Sciences (iCeMS), Kyoto University Institute for Advanced Study (KUIAS). This work was supported by a Grant-in-Aid for Scientific Research (no. 19H03136 to N.M.) and a Grant-in-Aid for JSPS Fellows (no. 19J23290 to S.H.) from the Japan Society for the Promotion of Science.

## Author contributions

The experiments were designed by S.H. and N.M., and were performed by S.H., with contributions from Y.K. and M.H. The manuscript was written by S.H. and N.M., with contributions from S.I. All authors read and approved the final manuscript.

---

**Supplementary Fig. 1.**
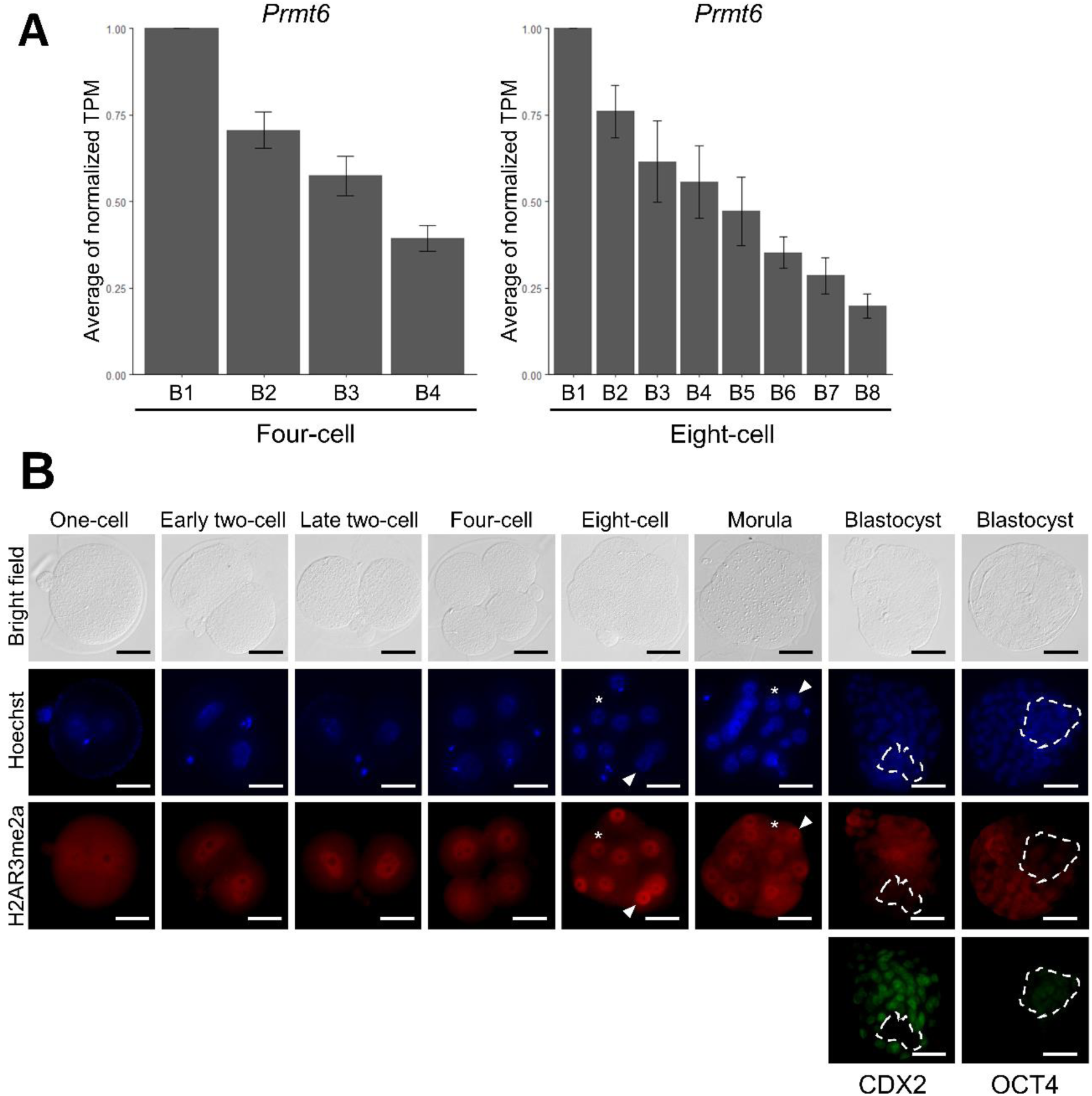
Asymmetric distribution of *Prmt6* mRNA and H2AR3me2a. (A) Average of the relative *Prmt6* mRNA levels in each blastomere at four- and eight-cell stage. Each blastomere at 4- or 8-cell stage is named B1 to B4 (4-cell, n=16) or B1 to B8 (8-cell, n=4) according to the amount of *Prmt6* expression. TPM (Transcripts Per Million) was normalized based on the blastomere with highest *Prmt6* expression in each embryo, and the mean averages are shown (± s.e.m.). Data were obtained from E-MTAB-3321. (B) Immunostaining for H2AR3me2a in embryos at each developmental stage. Hoechst (blue), H2AR3me2a (red), and CDX2 or OCT4 (green, only at the blastocyst stage) are shown. Scale bars, 50 μm. Arrowheads indicate the cell with high fluorescence and asterisks indicate the cell with low fluorescence in four- and eight-cell embryos. Dotted lines mark the inner cell mass in blastocysts.

**Supplementary Table 1.**
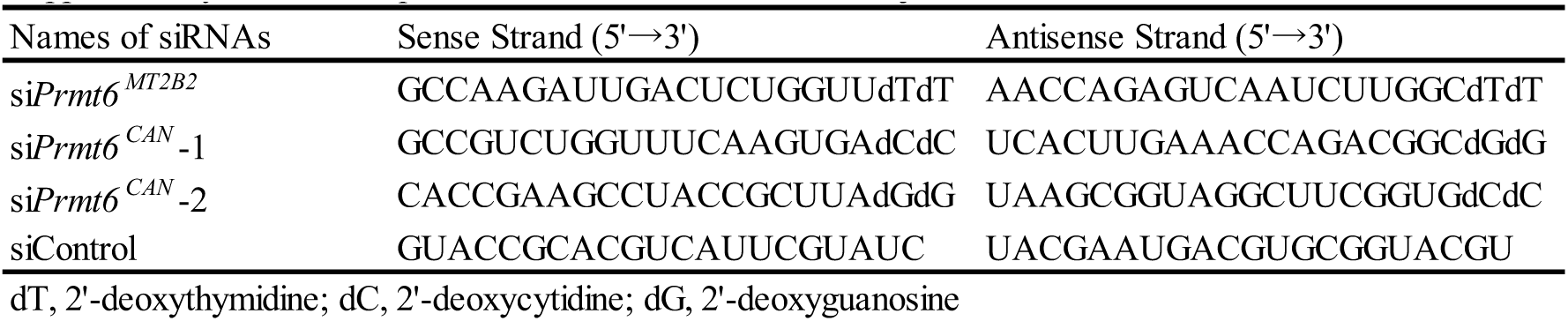
Sequences of siRNAs used for microinjection.

**Supplementarry Table 2.**
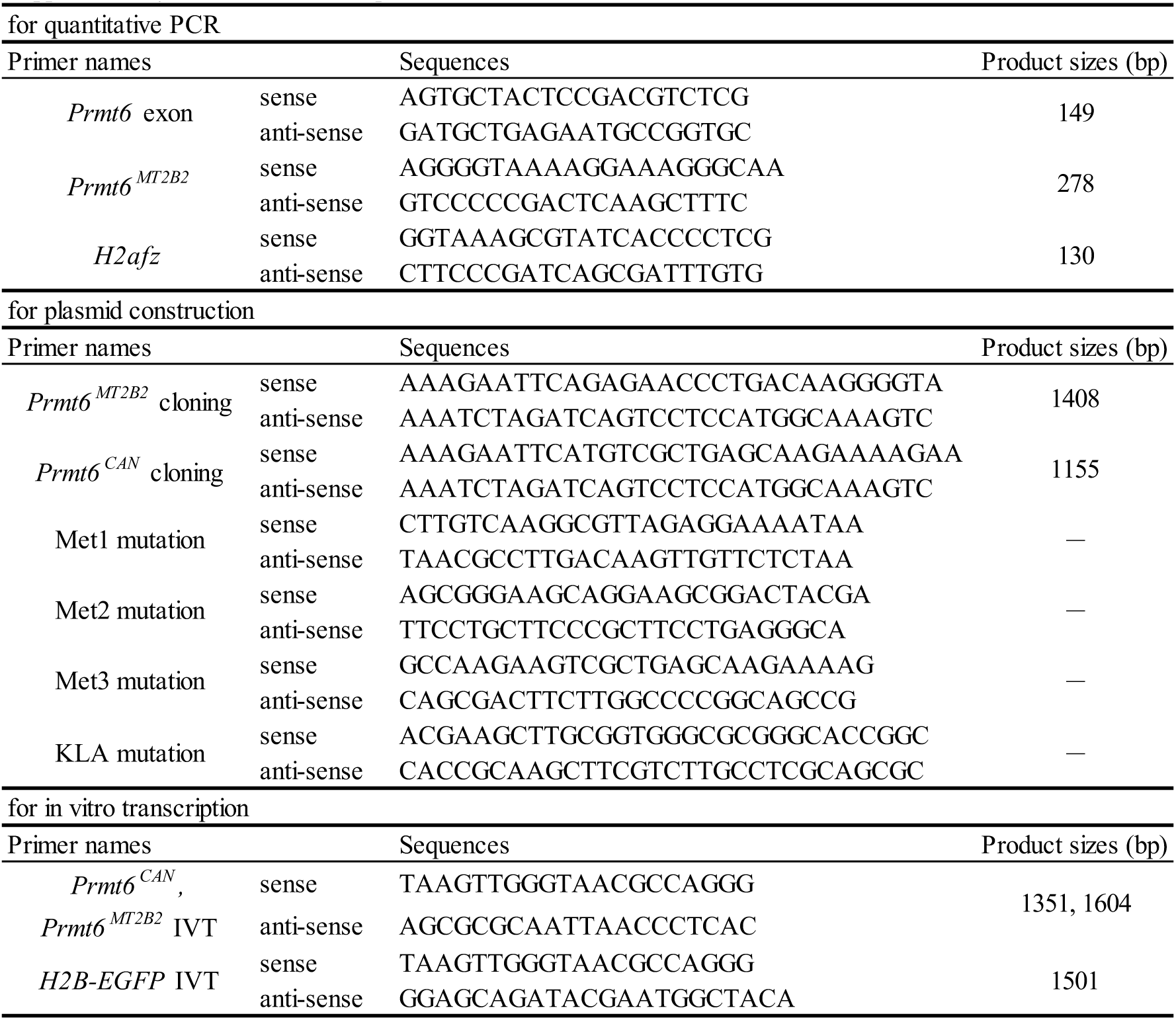
Primer sequences.

**Supplementary Table 3.**
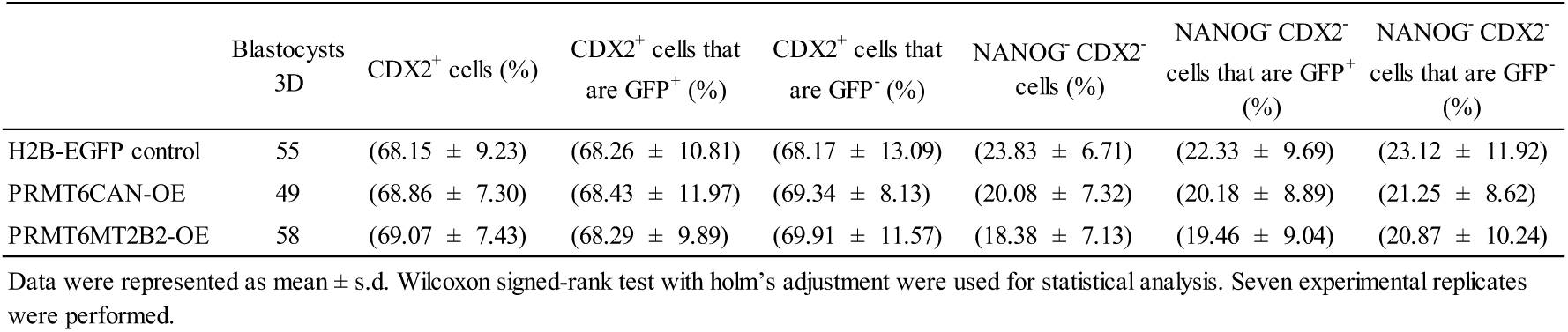
Analysis of the distribution of progeny of injected blastomere at the blastocyst stage based on Fig. 5B.

**Supplementary Table 4.** Analysis of the distribution of progeny of siRNA injected two-cell blastomere at the blastocyst stage.

|  | Blastocysts<br>3D | Total cells (n) | NANOG<sup>+</sup> cells<br>(%) | CDX2<sup>+</sup> cells (%) | GFP<sup>+</sup> cells (%) | NANOG<sup>+</sup> cells<br>that are GFP<sup>+</sup> (%) | NANOG<sup>+</sup> cells<br>that are GFP<sup>-</sup> (%) | GFP<sup>+</sup> cells that are<br>NANOG<sup>+</sup> (%) |
| --- | --- | --- | --- | --- | --- | --- | --- | --- |
| siControl | 37 | 77.62 ± 13.07 | (10.41 ± 7.91) | (73.01 ± 7.94) | (49.20 ± 7.24) | (11.17 ± 13.49) | (9.79 ± 8.35) | (49.36 ± 31.70) |
| siPrmt6-1 | 29 | 70.79 ± 12.34 | (11.74 ± 6.43) | (73.74 ± 7.99) | (44.19 ± 9.83)* | (14.40 ± 9.58) | (9.82 ± 7.47) | (54.58 ± 23.97) |
| siERVL-Prmt6 | 31 | 76.45 ± 12.63 | (10.49 ± 4.93) | (70.13 ± 11.13) | (48.20 ± 7.26) | (11.23 ± 7.31) | (9.65 ± 6.19) | (49.52 ± 27.86) |

Data were represented as mean ± s.d. Wilcoxon signed-rank test with holm's adjustment were used for statistical analysis, \*P < 0.05. Three experimental replicates were performed.

**Supplementary Table 5.** Analysis of the cell fate distribution of *EGFP, Prmt6 ^CAN^* or *Prmt6 ^MT2B2^* mRNA injected one-cell embryos at the blastocyst stage.

|  | Blastocysts | Total cells (n) | NANOG<sup>+</sup> cells (%) | CDX2<sup>+</sup> cells (%) | NANOG<sup>-</sup> CDX2<sup>-</sup> cells (%) |
| --- | --- | --- | --- | --- | --- |
| EGFP control | 39 | 77.49 ± 9.19 | (15.50 ± 5.51) | (77.26 ± 6.07) | (9.62 ± 6.63) |
| *Prmt6*<sup>CAN</sup>-OE | 35 | 78.11 ± 10.15 | (15.32 ± 6.98) | (73.85 ± 8.98) | (12.58 ± 7.54) |
| *Prmt6*<sup>MT2B2</sup>-OE | 34 | 78.21 ± 11.68 | (14.34 ± 4.64) | (73.31 ± 8.16)* | (13.22 ± 8.59)* |

Data were represented as mean ± s.d. Wilcoxon signed-rank test with holm's adjustment were used for statistical analysis, \*P < 0.05. Three experimental replicates were performed.

